# Prenatal diagnosis of *HNF1B*-associated renal cysts: Need to differentiate intragenic variants from 17q12 microdeletion syndrome?

**DOI:** 10.1101/576918

**Authors:** Georgia Vasileiou, Juliane Hoyer, Christian T. Thiel, Jan Schaefer, Maren Zapke, Mandy Krumbiegel, Cornelia Kraus, Markus Zweier, Steffen Uebe, Arif B. Ekici, Michael Schneider, Michael Wiesener, Anita Rauch, Florian Faschingbauer, André Reis, Christiane Zweier, Bernt Popp

## Abstract

**Objective:** Large deletions of chromosome 17q12 (17q12DS) or intragenic variants in *HNF1B* are associated with variable developmental, endocrine and renal anomalies, often already noted prenatally as hyperechogenic/cystic kidneys. Here, we describe pre- and postnatal phenotypes of seven individuals with *HNF1B* aberrations and compare their clinical and genetic data to previous studies.

**Methods:** Prenatal sequencing and postnatal chromosomal microarray analysis was performed in seven individuals with renal and/or neurodevelopmental phenotypes. We evaluated *HNF1B*-related clinical features from 82 studies and reclassified 192 reported intragenic *HNF1B* variants.

**Results:** In a prenatal case, we identified a novel in-frame deletion p.(Gly239del) within the *HNF1B* DNA binding domain, a mutational hotspot as demonstrated by spatial clustering analysis and high computational prediction scores. The six postnatally diagnosed individuals harbored 17q12 microdeletions of varying size. Literature screening revealed highly variable reporting of *HNF1B-*associated clinical traits. Overall, developmental delay was more frequent in 17q12DS carriers, although both mutation groups showed a high phenotypic heterogeneity. The reclassification of all previously reported intragenic *HNF1B* variants provided an up-to-date overview of the mutational spectrum.

**Conclusions:** We highlight the value of prenatal *HNF1B* screening in renal developmental diseases. Standardized clinical reporting and systematic classification of *HNF1B* variants is necessary for a more accurate risk quantification of pre- and postnatal clinical features, improving genetic counseling and prenatal decision-making.

## INTRODUCTION

The HNF1B (hepatocyte nuclear factor-1-beta, MIM *189907; also known as transcription factor-2 (*TCF2*)) protein is a transcription factor which, together with its dimerization partner HNF1A, belongs to the homeodomain-containing superfamily.[1] HNF1B is highly expressed in numerous fetal/adult tissues, where it mediates tissue-specific gene expression, development and function.[2, 3] It has a prominent role in fetal renal development, including regulation of organization and differentiation of renal epithelium,[1, 4, 5] urogenital formation,[4, 6] and tubular development in the nephron.[1]

Genetic variants affecting *HNF1B* are associated with multiple phenotypes.[7–11] Aberrations of *HNF1B* were initially described as causative for maturity-onset diabetes of the young type 5 (MODY5; MIM #137920).[1, 8, 11–13] Furthermore, *HNF1B* variants have been shown to be causal for the renal cysts and diabetes syndrome (RCAD; MIM #137920),[14] renal hypodysplasia (RHD),[15] familial glomerulocystic kidney disease (GCKD),[16] autosomal dominant tubulointerstitial kidney disease [ADTKD, also “medullary cystic kidney disease” (MCKD) or “juvenile hyperuricemic nephropathy” (FJHN)][17, 18] and congenital anomalies of kidney and urinary tract (CAKUT).[8, 11, 15, 16, 19] The latter is a frequent cause of severe fetal renal anomalies often leading to termination of pregnancy (TOP), neonatal death and chronic renal disease in children.[19–21]

Apart from diabetes mellitus and the highly heterogeneous renal phenotype, with the most common manifestations being hyperechogenic kidneys prenatally[22] and renal cysts postnatally,[11] *HNF1B* aberrations have been associated with several other clinical anomalies. These include agenesis or hypoplasia of the pancreas,[8, 19] impaired liver function,[23] urogenital anomalies,[24] electrolyte abnormalities[17, 25] and diaphragmatic hernia.[26]

*HNF1B* pathogenic variants include intragenic mutations such as single-nucleotide variants (SNVs), small insertions or deletions of bases (indels) and small copy number variants (CNVs) affecting only parts of the gene as well as larger CNVs encompassing the entire *HNF1B*.[7] SNVs/indels described to date include nonsense, frameshift, splice-site and missense variants. The latter are mainly distributed within exons 2-4, which code for the DNA-binding domain.[8, 19, 22, 27] All described *HNF1B* whole gene deletions so far, correspond to chromosome 17q12 microdeletions spanning 1,3 to 1,8 Mb on average and including neighboring genes.[10, 28, 29] Both, intragenic variants and genomic rearrangements contribute almost equally to the *HNF1B*-associated phenotype.[11]

Importantly, numerous studies reported neurodevelopmental disorders (NDDs) in individuals with 17q12 microdeletions (MIM #614527) containing *HNF1B.*[10, 28–33] Although NDDs were considered an exclusive feature of chromosome 17q12 microdeletion syndrome (17q12DS), there are recent reports also in individuals with intragenic *HNF1B* alterations.[16, 34–36]

In this study we report on a series of seven individuals with intragenic *HNF1B* aberrations or 17q12DS and describe their renal and extra-renal phenotype prenatally and in childhood or adult life. By comparing this cohort to reports from current literature, we discuss the potential need to differentiate intragenic variants from 17q12DS in the increasingly common setting of prenatally diagnosed kidney abnormalities. This question is not only important for prenatal decision-making but also for postnatal management in affected individuals.

## SUBJECTS AND METHODS

### Individual 1 (I1)

A pregnant woman presented to our Department of Obstetrics and Gynecology at gestational week 21 5/7 for sonographic examination. An isolated bilateral renal cortical hyperechogenicity without kidney enlargement was noted in the fetus. Amniotic fluid volume was normal, and amniocentesis was performed. Prenatal ultrasound at 32 weeks of gestation additionally revealed bilateral fetal pyelectasis. The female individual I1 was born at gestational week 36 6/7 with normal birth parameters. During the neonatal period she manifested a temporary respiratory adjustment disorder and feeding problems. Postnatal kidney ultrasound at 3 months of age demonstrated a regression of renal hyperechogenicity, bilateral renal cysts of varying size, with the largest being 3 mm, an enlarged right kidney with a total volume of 24 ml (97. P), renal pelvis dilatation grade I (8 mm) of the left kidney and bilateral megaureters of 2 mm. Sonographic examination of liver and pancreas exhibited no structural anomalies. Blood glucose level was slightly increased (110mg/dl), and urine albumin/creatinine ratio was 504mg/g. Plasma magnesium and uric acid concentration as well as liver transaminases were normal. Additional postnatal findings included a minor atrial septal defect. Ocular anomalies were not detected. At the time of last assessment at age of 3 months, development and growth were in the normal range. Renal sonographic examination of both parents did not reveal any abnormalities (Fig. 1C, 1D, Tab. 1 and Supplementary note).

**Fig. 1.**
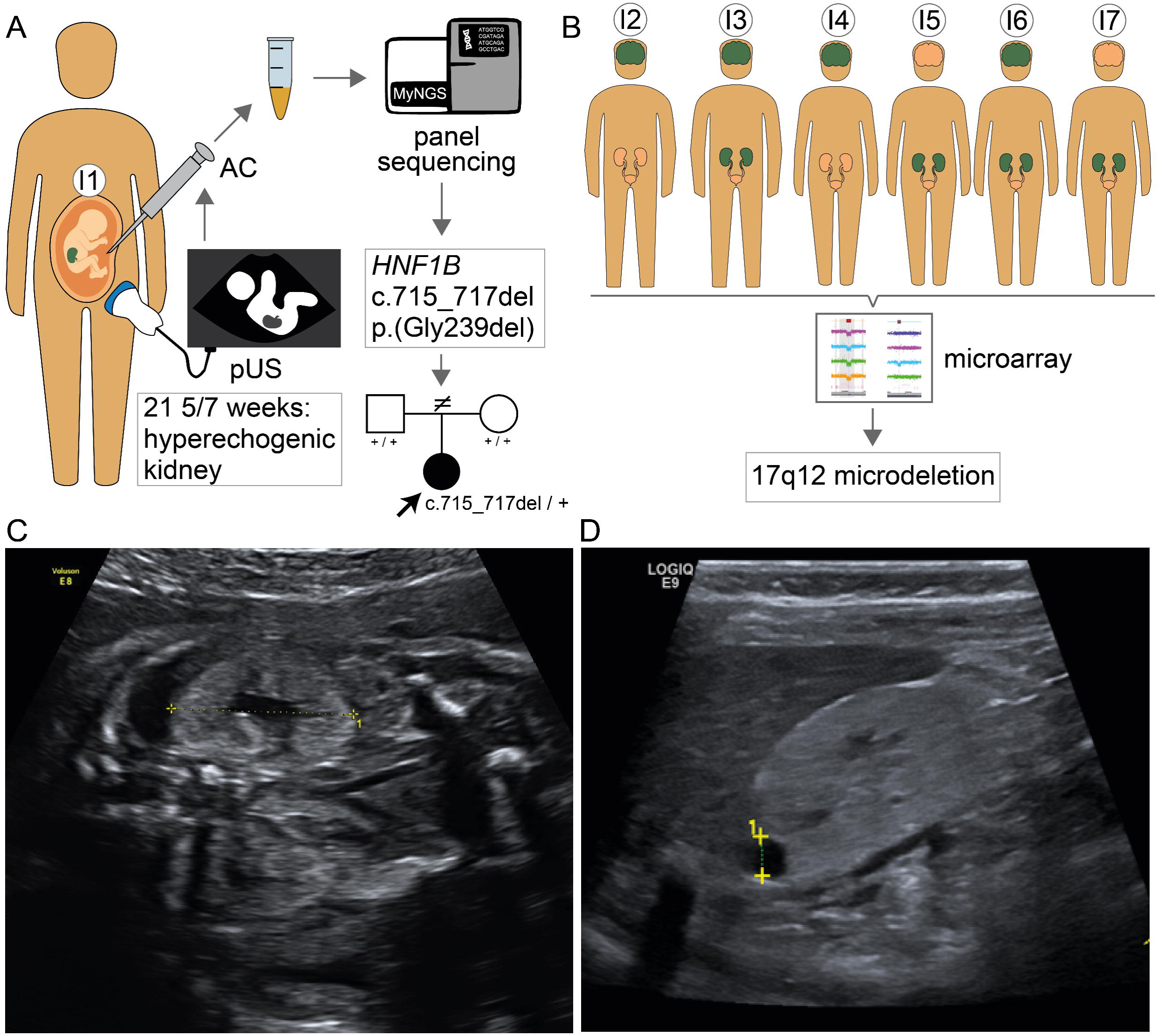
(A) Pictogram describing the prenatal diagnosis of hyperechogenic kidneys in individual I1 at gestational age 21 5/7 weeks followed by amniocentesis and targeted panel sequencing which identified the HNF1B variant c.715_717del. Postnatal segregation analysis in the healthy parents confirmed de novo occurrence of the variant (pedigree). (B) Pictogram summarizing the causes for referral of individuals I2-I7 with 17q12 microdeletions. Affected organ systems are marked in dark green: brain, neurodevelopmental disorder; kidneys, renal dysfunction; a combination thereof both. (C) Prenatal fetal sonography of individual I1 at gestational week 26 showing renal cortical hyperechogenicity with reduced corticomedullary differentiation. (D) Postnatal sonography of individual I1 at 5 months of age showing multiple small and one larger cyst of the right kidney.

**Tab. 1.** Clinical and molecular findings in 7 individuals with *HNF1B*-associated phenotype. The following abbreviations and symbols are used: +, present; -, absent; NA, not analyzed; ID, intellectual disability; IQ, intelligence quotient

|  | Individual 1 | Individual 2 | Individual 3 | Individual 4 | Individual 5 | Individual 6 | Individual 7<sup>[37]</sup> |
| --- | --- | --- | --- | --- | --- | --- | --- |
| **Causes of referral** | renal anomalies | neurodevelopmental disorder | neurodevelopmental disorder | neurodevelopmental disorder | renal anomalies | renal anomalies | renal anomalies |
| **Genetic change** | c.715-717del, p.(Gly239del) | 17q12del | 17q12del | 17q12del | 17q12del | 17q12del | 17q12del |
| **Inheritance** | *de novo* | *de novo* | Mother not carrier, father not available | *de novo* | *de novo* | *de novo* | *de novo* |
| **Diagnosis** | prenatal | postnatal | postnatal | postnatal | postnatal | postnatal | postnatal |
| **Sex** | female | male | male | female | female | female | female |
| **Birth parameters** | weight 2650 g<br>length 47 cm<br>head circ. 33 cm<br>Apgar 9/9/10 | weight 3620 g<br>length 53 cm<br>head circ. 36.5 cm<br>Apgar 10/10 | weight 3020 g<br>length 52 cm<br>head circ. 32 cm<br>Apgar 10/10 | weight 2730 g<br>length 48 cm<br>head circ. 35 cm<br>Apgar 10/10 | NA | NA | NA |
| **Age at latest follow-up** | 3 months | 8 years, 8 months | 15 years, 8 months | 12 years | 18 years | 28 years | 45 years |
| **Age at diagnosis of renal disease** | prenatal: 21 weeks, 5 days<br>postnatal: 3 months | - | 12-15 years | - | approx. 14 years | >26 years | 4 months |
| **Renal disease** | **prenatal:**<br>hyperechogenic kidneys,<br>pyelectasis<br>**postnatal:**<br>bilateral renal cysts<br>kidney enlargement<br>renal pelvis dilatation<br>bilateral megaureter | NA | bilateral renal cysts | - | bilateral renal cysts,<br>stage II of chronic kidney disease | recurrent urinary tract infections in childhood,<br>bilateral renal cysts,<br>end-stage renal disease | right kidney agenesis,<br>left kidney hypoplasia,<br>recurrent interstitial nephritis,<br>end-stage renal disease |
| **Developmental delay/**<br>**cognitive impairment** | - | learning difficulties | mild ID | motor delay in childhood,<br>learning and concentration difficulties,<br>low-normal IQ | - | motor delay in childhood | - |
| **Autism** | - | - | Asperger syndrome | autistic-like behavior | - | - | - |
| **Behavioral anomalies** | - | oppositional behavior | Hyperactivity,<br>Restlessness,<br>abnormal social behavior | - | - | - | - |
| **Epilepsy** | - | - | - | - | - | single seizure attack in adulthood | - |
| **Brain anomalies** | NA | - | NA | enlarged brain ventricles | - | NA | NA |
| **Diabetes** | 110mg/dl | NA | - | - | + | + | - |
| **Other extra-renal manifestations** | atrial septal defect | ventricle septum defect | - | short stature,<br>increased hepatic enzymes<br>Brown's syndrome | mild strabism | increased hepatic enzymes,<br>biliary cirrhosis due to liver cysts | strabism convergens,<br>uterus aplasia, ovarian cysts, aortic insufficiency,<br>joint hyperextension |
| Abnormal magnesium concentration | - | NA | - | NA | NA | NA | NA |
| Hyperuricemia | - | NA | NA | NA | + | NA | NA |

### Individuals 2-7 (I2-I7)

Clinical data of individuals I2-I7, who presented with renal and/or neurodevelopmental phenotypes at our genetic outpatient clinic and at the Center for Rare Diseases Erlangen, were retrospectively collected for this study (Fig. 1B). Detailed clinical descriptions are provided in Tab. 1 and Supplementary note. Individual I7 was previously described.[37]

### Genetic analyses including sequencing and chromosomal microarray

Details for next-generation sequencing (NGS) based panel and Sanger sequencing methods as well as chromosomal microarray analysis (CMA) methods are described in Supplementary note and Supplementary File 2. In brief, a custom panel was sequenced on a Illumina MiSeq, processed bioinformatically as described [38] and analyzed for variants in four genes (*PKD1, PKD2, PKHD1, HNF1B*) associated with polycystic kidney disease. Sanger sequencing was used to confirm and validate the segregation of identified variants and Long Range PCR with subsequent Sanger sequencing was used to analyze the duplicated regions of *PKD1*. CMA data was analyzed for CNVs ≥ 100 kb using Software ChAS and evaluated against control databases (Supplementary note).[39]

### Computational analyses of ***HNF1B* variants**

The clustering analysis of described *HNF1B* variants, the collection of these variants from the literature together with computational analysis of all possible single amino acid deletions and missense substitutions as well as the protein structure analysis of the Gly239del variant are described in the Supplementary note and all data is provided in Supplementary File 1. In brief, we collected an up-to-date list of 217 intragenic *HNF1B* variants from 88 articles and public databases, harmonized these according to the HGVS nomenclature, annotated them with different computational scores and evaluated their pathogenicity according to the ACMG criteria. To analyze the spatial clustering, we plotted the distribution of these variants across the linear protein representation and estimated p-values from empiric distributions of drawing the observed number of missense variants in the respective domains. Additionally, we used the published tertiary protein structure of *HNF1B* to analyze the potential effect and proximity to other pathogenic variants of the herein identified c.715_717del, p.(Gly239del) variant.

## RESULTS

### Prenatal identification of a pathogenic in-frame deletion in individual I1

Prenatal panel sequencing in individual I1 revealed a novel heterozygous 3 base pair (bp) deletion in exon 3 of *HNF1B* (NM_000458.3: c.715_717del), resulting in an in-frame deletion of the highly conserved amino acid (AA) glycine at position 239 (p.(Gly239del)) (Fig. 2B). The identified indel variant was not present in the Genome Aggregation Database (gnomAD) and was computationally predicted (CADD, MutationTaster) to be deleterious. A potential splice effect was not detected (Human Splicing Finder, version 3.1). The c.715_717del variant could not be identified in DNA from peripheral blood lymphocytes of both parents and thus likely arose *de novo.* Structural modeling based on the crystal structure of the HNF1B protein showed that the glycine at position 239 (Gly239) lies in an alpha-helix motif of the homeodomain, which has DNA binding function. The deletion of Gly is predicted to disrupt two neighboring AAs (Trp238 and Lys237) involved in binding to DNA (Fig. 3). Two missense variants (c.715G>C, p.(Gly239Arg)[40]; c.716G>A, p.(Gly239Glu)[41]) affecting the same AA were reported in a boy with bilateral renal cysts, kidney failure and diabetes in childhood and in a girl with MODY, renal cysts in right kidney, agenesis of left kidney and pancreas atrophy, respectively. Furthermore, a variant affecting the neighboring residue Trp238 (c.712T>C, p.(Trp238Arg)), was described as pathogenic (Fig. 3).[9] Based on the ACMG criteria[42] (*de novo* occurrence, no inclusion in population databases, previously described variant at the same position, affected highly conserved AA, computationally indicated pathogenicity), we classified the identified variant as pathogenic (class 5).

**Fig. 2.**
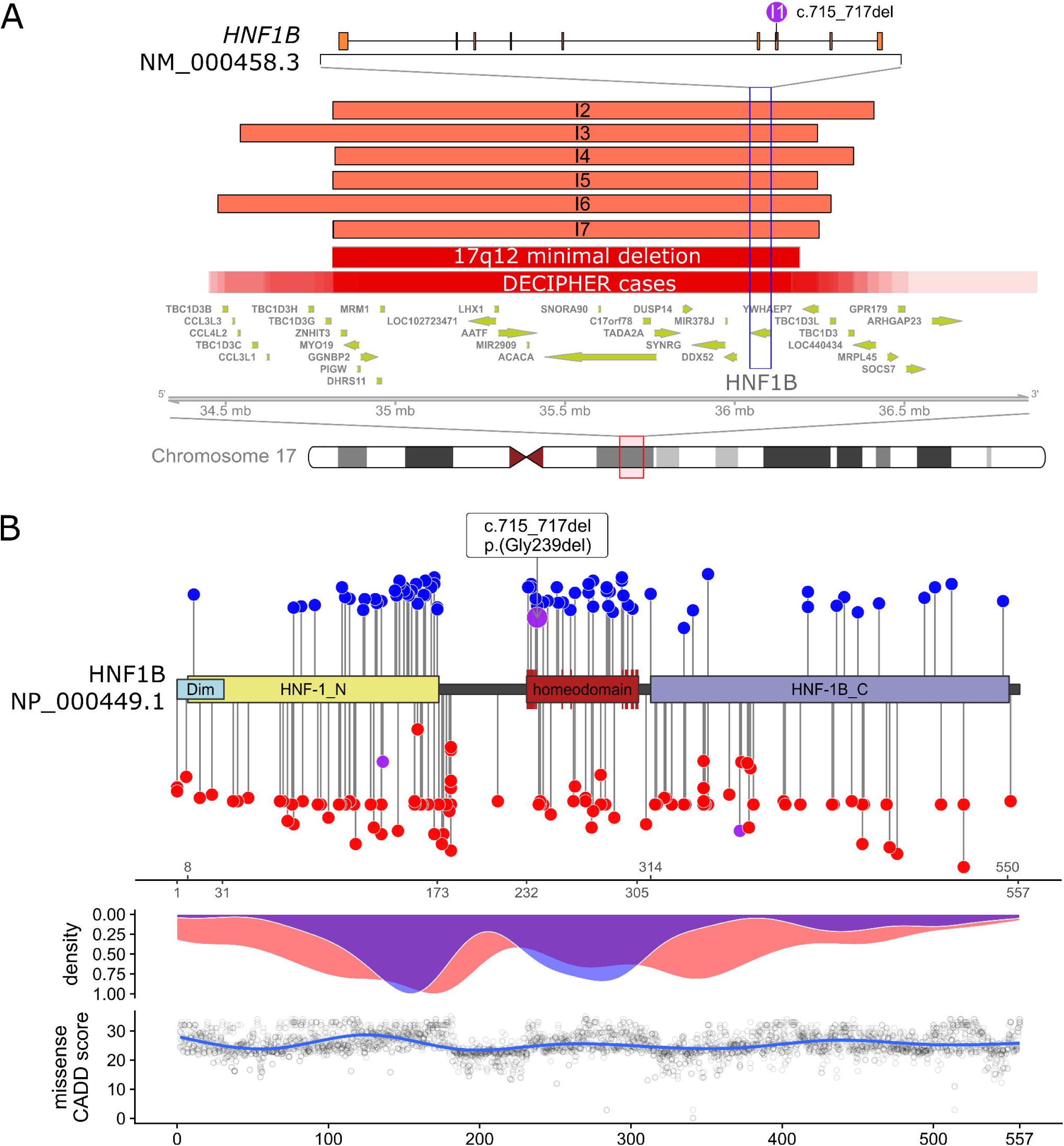
(A) 17q12 microdeletions on chromosome 17q12 (red box) identified in individuals I2-I7 (light red labeled bars) with location of genes typically affected by these CNVs (see also Supplementary File 1, sheet “CNV_genes”). Dark red bars indicate the minimal 17q12 microdeletion and the variability observed in 33 cases from the DECIPHER database, respectively. The location of *HNF1B* gene within the CNV is highlighted (blue box) and the exon structure for transcript NM_000458.3 is indicated above the blue box. Location of the c.715_717del variant in exon 3 is indicated by the purple circle (B) Upper panel: Linear schematic representation of the HNF1B protein (based on NP_000449.1) with domains (limits indicated by ticks below x-axis) and localization of all described (likely) pathogenic variants (length of the segments corresponds to the variants CADD score). The small red segments on the homeodomain represent AA residues directly involved in DNA binding or with specific DNA base contacts. Dim, N-terminal dimerization domain specific for the formation of homo- or heterodimers with HNF1A; HNF-1_N, N-terminal domain involved in DNA binding; homeodomain, DNA binding domain; HNF-1B_C, C-terminal transactivation domain which mediates the transcription and recruitment of coactivators.[57] The blue circles indicate all missense variants (above the schematic) and the red all truncating variants (below the schematic) described in the literature and databases. The in-frame indel c.715_717del, p.(Gly239del) (above the schematic and presented as larger circle) and two in-frame deletions of whole exons (below the schematic) are presented in purple. Middle panel: Density plot of truncating (red) and missense (blue) variants reported. Missense variants show a maximal density at AA 154 in the HNF-1_N domain and a local maximum at AA 280 in the homeodomain. Lower panel: Generalized linear model of the CADD score for all possible missense variants (circles) in *HNF1B*.

**Fig. 3.**
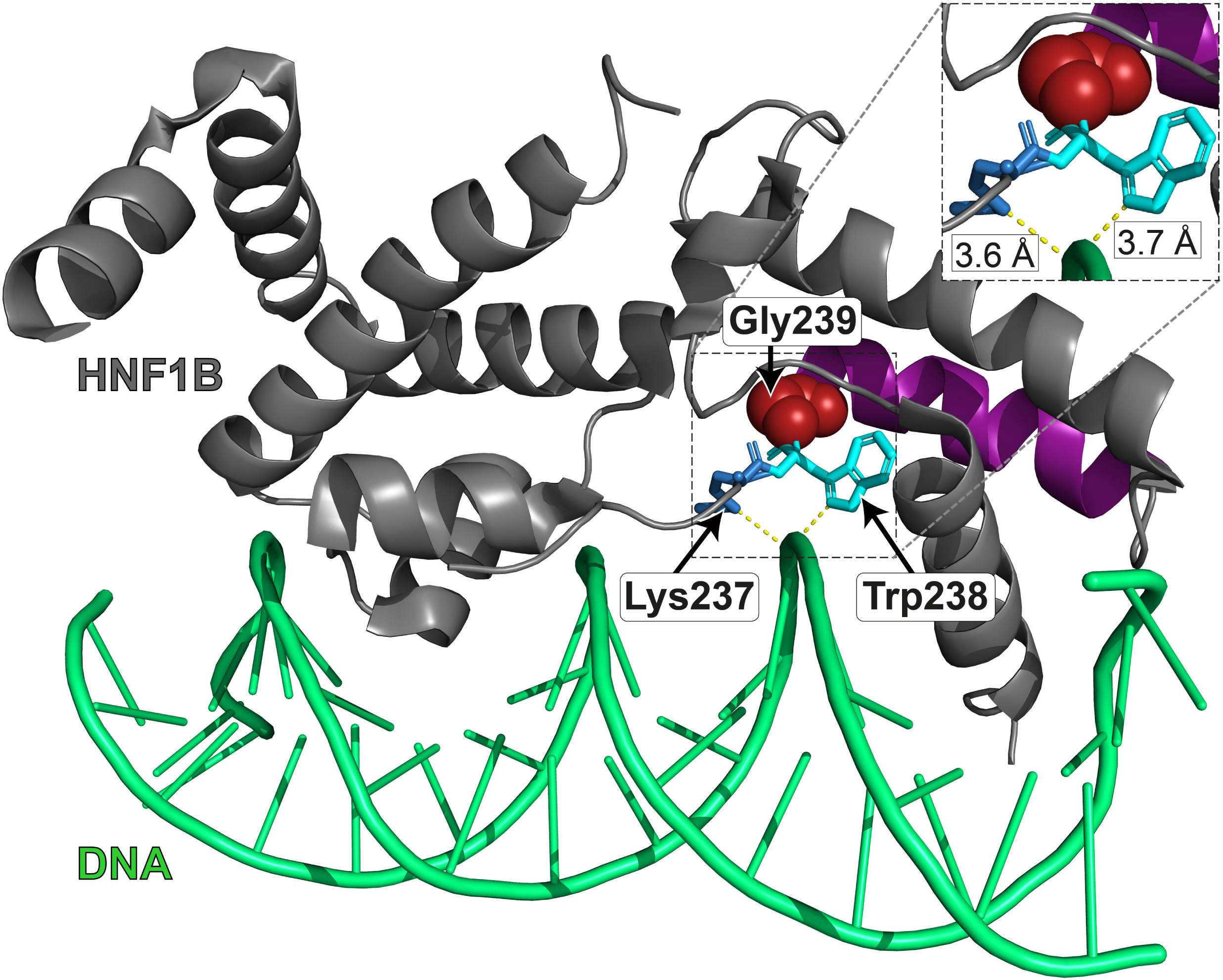
Crystal structure of HNF1B (grey) monomer bound to the DNA double helix (green) based on PDB 2H8R[57]. The AA residue Gly239 mutated in individual I1 is shown in sphere representation (red). The neighboring residues Trp238 (cyan) and Lys237 (blue) are annotated as having a “specific DNA base contact[s]” or being involved in “DNA binding”, respectively (based on NCBI NP_000449.1). Additionally, the variant c.712T>C, p.(Trp238Arg) has been reported as pathogenic.[9] The typical alpha helix breaker glycine at position 239 is located at the end of an alpha-helix motif (residues 240-252; purple) and its deletion (I1 described herein) or substitution[40, 41] likely disrupt the conformation and positioning of the AAs Lys237 and Trp238 required for DNA binding of the HNF1B transcription factor. The dashed box shows a close-up view affected region and the distances between the AA residues and the DNA double helix in angstrom.

### 17q12 microdeletions in individuals I2-I7

Postnatal high-resolution CMA analysis in individuals I2-I7 revealed 17q12 microdeletions of variable sizes ranging from 1.42 Mb in I5 to 1.80 Mb in I6 (Fig. 2A). *De novo* occurrence was confirmed in all cases apart from individual I3, where the deletion was excluded in the mother, but paternal DNA was not available (see also Tab. 1 and Supplementary note).

### Clinical features of the affected individuals

Prenatal renal anomalies such as hyperechogenic kidneys were only identified in individual I1 with the in-frame deletion in *HNF1B* (Fig. 1C). Postnatally, 4/7 individuals exhibited renal cysts. Individual I1 was diagnosed with multiple renal cysts in early infancy (Fig. 1D), I3 and I5 in puberty and I6 in adulthood. Other renal abnormalities included kidney enlargement, renal pelvis dilation and bilateral megaureter in individual I1 as well as right kidney aplasia and left kidney hypoplasia in individual I7. Recurrent urinary tract or kidney infections from childhood on were reported in individuals I6 and I7. Individuals I5 and I7 developed stage II and end-stage renal disease in their late teens and I6 end-stage renal disease after the age of 26. In individuals I2 and I4 no renal manifestations were reported at last follow-up, at the age of 8 years and 8 months, and 12 years, respectively.

Two individuals had diabetes, both diagnosed in puberty (I5 and I6). I1 showed increased blood glucose levels at 3 months of age. Other extra-renal manifestations included strabism (I4, I5 and I7), elevated liver enzyme levels (I4 and I6), liver cysts and primary biliary cirrhosis (I6), hyperuricemia (I5), short stature (I4), uterus aplasia and ovarian cysts (I7) and cardiac anomalies (I1, I2 and I7).

The most prominent extra-renal phenotypes in individuals with 17q12DS were NDDs and behavioral problems (Fig. 1B). Individual I2 exhibited learning difficulties, I3 mild intellectual disability and Asperger syndrome and I4 a low-normal IQ with learning and concentration difficulties. Behavioral problems varied from oppositional behavior (I2) and hyperactivity and restlessness (I3) to autistic-like behavior (I4). Motor milestones were delayed in I4 and I6. Individual I6 had a single seizure attack in adulthood. Finally, MRI showed enlarged brain ventricles in individual I4 (Tab. 1 and Supplementary note).

### Reviewed intragenic variants and clustering in protein domains

We retrieved a total of 217 intragenic *HNF1B* variants from databases, 88 published articles and this report (I1) (Supplementary File 1). Only 46/216 variants (21.3%; excluding the variant in individual I1) were deposited in public databases as of 2018-12-02. After manual review, 192 of them could be classified as ACMG class 4 (likely pathogenic; n=57) or 5 (pathogenic; n=135). For four variants described in the literature there was enough evidence to classify them as ACMG class 2 (likely benign), while for 21 variants only insufficient information was available, and they were classified as ACMG class 3 (uncertain significance). The majority of (likely) pathogenic variants in *HNF1B* are truncating (59.4%; 114/194), whereas missense variants, clustering in important protein domains, constitute the second largest group (39.1%; 75/194) (Fig. 2B and Supplementary File 1).

Besides the c.715_717del variant in individual I1, two larger indels of 30 (c.1118_1147del) and 75 (c.410_484del) bp are annotated as in-frame variants deleting 10 (p.(Ala373_Gln383delinsGlu)) or 25 (p.(Arg137_Lys161del)) AAs, respectively. While these two variants are annotated as “disruptive” (Fig. 2B, marked in purple below protein scheme) because they ablate multiple AAs which likely disrupts the protein structure, the herein identified in-frame deletion of glycine at position 239 (p.(Gly239del)) is annotated as “conservative” (Fig. 2B, marked in purple above protein scheme). Besides affecting an AA position previously described as mutated in patients with HNF1B-associated disease, the deletion is located within the HNF1B homeodomain, which mediates DNA binding.

The HNF1B homeodomain and the second half of the N-terminal domain (“HNF-1_N”) host the two mutational hotspots for missense variants, while likely gene disruptive variants are dispersed throughout the protein (Fig. 2B, middle panel). The possible missense variants in these two domains additionally have significantly higher CADD scores when compared to those of the C-terminal domain (“HNF-1B_C”) or non-domain regions of the HNF1B protein (Fig. 2B, lower panel and Fig. S1).

## DISCUSSION

Improved sonographic technologies increasingly enable the identification of fetal hyperechogenic and/or cystic kidneys. Nevertheless, identifying the underlying cause and specifying prognosis remain challenging. The recurrent 17q12 microdeletion encompassing *HNF1B* constitutes a frequent differential diagnosis for fetal renal anomalies with variable postnatal consequences ranging from isolated renal to more complex, syndromic manifestations.[11, 15, 19, 22] The emerging role of *HNF1B* intragenic variants for prenatally detected renal and extra-renal phenotypes has only recently been recognized.[11]

Panel sequencing of common genetic causes for prenatal-onset kidney anomalies in individual I1 with hyperechogenic kidneys in fetal ultrasound, identified the novel in-frame single AA deletion c.715_717del, p.(Gly239del) in *HNF1B.* This is the first report of an in-frame *HNF1B* variant leading to a renal phenotype resembling that of previously described *HNF1B* cases. One of the main questions in this prenatal situation was further prognosis. Due to the variable presentation of *HNF1B* aberrations with renal and extra-renal plus possible neurodevelopmental aspects, this is particularly challenging. Currently prenatal prognosis and decision-making regarding TOP depend mainly on clinical presentation including oligohydramnios and detectable congenital anomalies. However, other issues to consider during prenatal genetic counseling without severe complications are frequency and severity of the various postnatal manifestations.

In this regard, the association of 17q12DS with a wide spectrum of NDDs in 30-89% individuals is of particular importance.[10, 29–33, 43] In our cohort, three individuals diagnosed with 17q12DS (I2-I4) presented primarily with psychomotor deficits and behavior anomalies, and two (I4 and I6) required therapy for delayed motor development in childhood. Although very young, individual I1 with the intragenic *HNF1B* variant did not show any signs of developmental delay. Clissold and colleagues provided evidence for association of NDDs exclusively with 17q12 microdeletions but not with *HNF1B* intragenic variants.[30] The predominant hypothesis for the neurodevelopmental phenotypes was haploinsufficiency of other genes encompassed by the deletion.[28, 29, 31, 33, 43] Until recently, only single cases of NDDs in carriers of intragenic *HNF1B* variants had been reported.[16, 34, 36] However, a recent study described NDDs in (6/54) 11% of individuals with MODY harboring *HNF1B* intragenic alterations; this frequency was significantly higher when compared with cases who had another cause of diabetes and not significantly different from 17q12DS cases (17.0%; 9/53).[35] This study only excluded CNVs and Fragile X syndrome in individuals with NDDs without further application of exome sequencing. This leaves the possibility of a second, independent variant causing a blended phenotype, as has been shown for 4.9% of the cases in a large cohort of individuals receiving exome sequencing for different diseases.[44] After the report of NDDs in carriers of intragenic variants, *HNF1B* has been included in curated gene lists for NDDs (SysID[45]; PanelApp “Intellectual disability Version 2.595”), without experimental data supporting the clinical observations. Contradicting this association, no intragenic *HNF1B* variant has been observed in several thousand individuals of large cohorts with autism[46] or developmental disorders.[47] Thus, it still remains unclear whether there is a causal link of *HNF1B* loss-of-function to NDDs. The difficulty of quantifying the NDD risk in both groups with *HNF1B* deletions and intragenic alterations is a primary cause of parental dilemma during prenatal decision-making. Further complicating estimation, published patient cohorts and case reports are often limited to describing only a subset of *HNF1B*-associated clinical traits, as shown by our evaluation for 12 *HNF1B*-associated phenotypes mentioned in 82 published studies reporting phenotype data on born individuals (Tab. 2; Supplementary File 1). New studies should follow a minimal, standardized clinical reporting scheme (Tab. 2) to enable fast systematic reviews and facilitate future estimation of prevalence for important symptoms like NDDs. Next to this clinical evaluation, behavioral analysis experiments of *HNF1B* knockout animal models may provide further insight of the involvement of this gene in neurodevelopment.

**Tab. 2.** Reported prevalence for 12 clinical features associated with 17q12DS or *HNF1B* variants and frequency of these phenotypes as described in 82 published articles with clinical data on born individuals. HPO, Human Phenotype Ontology terms; *, fraction of publications mentioning the respective feature (e.g. absence or presence count as mentioned)

| Phenotype | HPO | Reported* | Prevalence in literature |
| --- | --- | --- | --- |
| Abnormality of the kidney | HP:0000077 | 76/82 (92.7%) | 61%- 91% |
| Neurodevelopmental delay | HP:0012758 | 10/82 (12.2%) | 30%-89%: 17q12 deletions<br>11%: *HNF1B* variants |
| Diabetes mellitus | HP:0000819 | 76/82 (92.7%) | 9%-82% |
| Abnormality of the genital system | HP:0000078 | 39/82 (47.6%) | 18%-80% |
| Elevated hepatic transaminase | HP:0002910 | 35/82 (42.7%) | 12,5%-84% |
| Abnormality of the liver | HP:0410042 | 24/82 (29.3%) | 20%-32% |
| Abnormality of the pancreas | HP:0001732 | 33/82 (40.2%) | 6%-54% |
| Abnormality of brain morphology | HP:0012443 | 3/82 (3.7%) | 4%-50% |
| Short stature | HP:0004322 | 16/82 (19.5%) | 4% |
| Abnormality of the eye | HP:0000478 | 7/82 (8.5%) | 40%-44% |
| Abnormal magnesium concentration | HP:0004921 | 21/82 (25.6%) | 25%-75% |
| Hyperuricemia | HP:0002149 | 31/82 (37.8%) | <10-67% |

Postnatally, the presence of accompanying organ anomalies is important to guide appropriate management and surveillance. In the present study 5/7 (71.4%) individuals with either *HFN1B* intragenic variant or larger deletion had renal manifestations, in agreement with the previously reported 61-91% prevalence in *HNF1B* cases (Tab. 2).[9, 48] Absence of renal disease in individuals I2 and I4 may be attributed to a milder renal phenotype overshadowed by developmental deficits[33] or to their young age. Even though renal anomalies were only prenatally reported in individual I1 in our cohort, we cannot exclude that they were present but not detected in the remaining cases with renal manifestations identified after birth (I3 and I5-I7). In the literature, the severity of prenatal renal phenotypes is not correlated with the type of *HNF1B* aberrations.[11] Notably, *HNF1B* intragenic variants progressively result in more severe renal function impairment and poorer renal prognosis postnatally than 17q12 deletions.[9, 30] Accordingly, renal cysts were diagnosed in individual I1 directly after birth, whereas 4/6 (66,7%) 17q12DS individuals showed renal abnormalities later in infancy, puberty or early adult life. Regardless of the initial clinical indication for testing, identification of 17q12 deletions should always be accompanied by regular blood and urine tests, and sonographic kidney evaluation.

The frequency of diabetes in individuals with *HNF1B*-related clinical phenotypes is highly variable, ranging from 9-82% with a mean-age of diagnosis at 24 years, but also possible neonatal onset (Tab. 2).[9, 12, 48, 49] Consistently, individual I1 showed slightly increased levels of blood glucose in early infancy, whereas two 17q12DS individuals (I5 and I6) were diagnosed in puberty. The remaining individuals did not exhibit diabetes to date, yet at least for individuals I2-I4 this may relate to their young age. The severity of diabetes was shown to be similar in both carriers of truncating and missense variants. Dubois-Laforgue et al. reported a lower BMI and more severe diabetes observed at diagnosis in 17q12DS as compared to intragenic variant cases.[9] In addition to annual HbA1c and glucose level measurements both, carriers of 17q12DS and intragenic *HNF1B* variants, should be trained to self-monitor diabetes symptoms in order to enable early diagnosis and avoid complications.

Other noteworthy clinical features of the herein described individuals are short stature in individual I4, uterus aplasia in individual I7 and strabism in individuals I4, I5 and I7. Gonadotropin treatment increased the predicted final adult height of individual I4 for about 10 cm. Yet, reports on growth restriction in individuals with 17q12DS[29] or *HNF1B* intragenic variants[36] are rare, and the efficacy of growth hormone treatment in these cases needs elucidation. Uterus aplasia is within the *HNF1B*-associated spectrum, with intragenic variants and whole deletions reported in 18-50% of women with uterine and renal anomalies.[9, 24] Thus, obstetric evaluation already at young age is required. Finally, detailed ophthalmic examination is warranted, since in 40-44% of *HNF1B*-aberration carriers strabism and other ocular anomalies were reported (Tab. 2).[10]

The reasons for the phenotypic variability within the two groups of individuals with *HNF1B* intragenic alterations and whole deletions, as well as between these groups, currently remain unclear. The clinical diversity observed within the 17q12DS carriers could be attributed to the incomplete penetrance and variable expressivity of this CNV, similar to other recurrent microdeletions.[50] A quantitative impact of the 17q12 microdeletion with a relatively small size effect, which is difficult to be ascertained from the small cohorts examined, could for instance justify the presence or absence of NDDs in affected individuals.[50, 51] Regarding clinical differences within the intragenic variants group, different pathomechanisms, especially for the missense variants, constitute a possible hypothesis. Indeed, a dominant-negative effect, allowing a residual function of the HNF1B protein, rather than haploinsufficiency has been discussed for some variants.[52] The differential phenotypic presentation between the 17q12DS and intragenic aberration cohorts could be explained by the concurrent loss of other genes encompassed in the microdeletion, and interruption of interaction cascades with transcription factors or other regulatory elements. For instance, haploinsufficiency of two other genes (*ACACA*, MIM *200350 and *ZNHIT3*, MIM *604500) was indicated as the cause of the leaner figure and more severe diabetes at onset in 17q12DS cases.[9] This hypothesis was also suspected for the presence of NDDs in individuals with microdeletions. [33, 43] From the remaining 37 protein-coding genes encompassed in the deletion 7 have been implicated with NDDs, of which only *PCGF2* (MIM *600346) with dominant[53] and *PIGW* (MIM *610275) with recessive inheritance[54] have a confirmed association, whereas the rest are considered candidate genes. Consequently, *PCGF2* is the most notable candidate, however, it does not seem to be intolerant to heterozygous loss-of-function and missense variants (pLI=0.85, Z-score=1.21), and all described *PCGF2* carriers have the same recurrent missense variant with dominant-negative effect (Supplementary File 1).[53] In agreement, in the largest study attempting a genotype/phenotype correlation in 75 individuals with *HNF1B* alterations, no clear phenotypic differences regarding renal disease were identified between 17q12DS and intragenic variant carriers.[11] Therefore, the possibility remains that *HNF1B* is indeed the critical gene for all associated phenotypes, and that most of the variability is caused by additionally modifying genetic and developmental factors.

To date, only one recent study applying exome sequencing for CAKUT cases prenatally detected a *HNF1B* frameshift variant.[55] Likewise, the herein reported prenatal identification of an intragenic *HNF1B* variant in individual I1 is likely a consequence of increased availability and demand for such next-generation sequencing based techniques. Due to the known high frequency of intragenic *HNF1B* variants in fetal autopsy cases[8, 19, 27, 56] or in postnatally diagnosed children with renal diseases[11, 15, 22, 30, 34, 36], *HNF1B* mutation screening should be an integral part of prenatal diagnosis for hyperechogenic/multicystic kidneys. Customized targeted panels capturing multiple cystic kidney disease-related genes, like the one used herein, have several favorable features for the prenatal setting: high coverage, short turn-around times and compatibility with small benchtop sequencing machines. Clinical exome sequencing (covers ∼5.000 to 7.000 disease genes), is currently a reasonable compromise, although it requires more time and is less accessible. Regardless, NGS should be recommended concurrently with CMA for rapid prenatal diagnosis in fetuses with hyperechogenic or cystic kidneys. However, the application of next-generation methods will consequently result in the increased identification of pathogenic variants and variants of unknown significance (VUS). To facilitate fast and effective assessments, all identified variants should be systematically classified and deposited in appropriate databases, as performed in this study for all described in the literature so far (Supplementary note and Supplementary File 1).

In conclusion, similar to previous studies our study indicated no differences in the prenatal renal phenotypical spectrum between intragenic *HNF1B* aberrations and 17q12DS, and a wide range of postnatal renal and extra-renal abnormalities for both. The most prominent postnatal phenotypic differences, potentially affecting prenatal decision-making, are a more severe progress of renal impairment in individuals with intragenic *HNF1B* variants, and the increased NDD risk, which is a confirmed feature in 17q12DS and suspected in *HNF1B* intragenic variant carriers. Incomplete references to the overall clinical presentation of the affected individuals in the published reports and previous classifications of identified variants not following current recommendations of a standardized 5-tier system add to the difficulty of risk estimation. We therefore systematically collected *HNF1B*-related clinical traits, revised and harmonized all intragenic variants from 88 articles and propose a minimal reporting scheme for *HNF1B*-associated disease (Tab. 2). Building on these efforts, future studies using a standardized ascertainment of phenotype/genotype information will improve our understanding and characterization of *HNF1B*-associated disorders, and enable the optimization of genetic counseling.

## Authors’ contributions

B.P. and G.V. conceived the study. B.P., J.H., C.T.T., J.S., M.Za., M.S., F.F. and C.Z. provided patients’ data and performed clinical assessments. B.P., M.K., C.K., S.U., A.B.E., and A.Re. analyzed and interpreted the molecular data. M.Zw. and A.Ra. designed and provided the custom panel. B.P. created figures and Supplementary files. B.P. and G.V. performed the variant review and standardization. G.V. and C.Z. retrospectively collected patient data and wrote the case reports. G.V., C.Z. and B.P. wrote and edited the manuscript. J.H., C.T.T., J.S., M.Za., M.K., C.K., M.Zw. S.U., A.B.E., M.W.B, M.W., A.Ra. F.F., and A.Re. reviewed the draft manuscript.

## Compliance with ethical standards

### Ethical aspects

Informed written consent was obtained from all patients or their legal guardians, and the study was approved by the Ethical review board of the Friedrich-Alexander-Universität Erlangen-Nürnberg.

## WEB LINKS

CADD: https://cadd.gs.washington.edu/score/

gnomAD: http://gnomad.broadinstitute.org/

MutationTaster: http://www.mutationtaster.org/

SnpEff/SnpSift: http://snpeff.sourceforge.net/

DECIPHER: https://decipher.sanger.ac.uk/

dbNSFP: https://sites.google.com/site/jpopgen/dbNSFP/

Genomics England PanelApp: https://panelapp.genomicsengland.co.uk/

## WHAT’S ALREADY KNOWN ABOUT THIS TOPIC

- *HNF1B* intragenic variants and recurrent deletions of the 17q12 chromosomal region are correlated with hyperechogenic/cystic kidneys prenatally and renal anomalies postnatally
- Psychomotor delay is an additional manifestation of *HNF1B*-phenotype primarily associated with 17q12 microdeletions
- Currently chromosomal microarray, but not *HNF1B* sequencing, is common in prenatal diagnostics of fetuses with renal anomalies
- Incomplete clinical descriptions and variant classifications in the literature complicate risk estimation

## WHAT DOES THIS STUDY ADD?

- We describe a novel prenatally identified in-frame deletion in *HNF1B* in a fetus with hyperechogenic/cystic kidneys
- We aim at raising the attention of clinicians to the value of often overlooked prenatal *HNF1B* screening using next-generation sequencing
- We systematically collected clinical features from 82 studies with *HNF1B* aberrations to improve understanding and prognosis assessments
- We re-classified 192 variants based on ACMG criteria to facilitate fast and effective diagnosis by clinicians and future studies

## SUPPORTING INFORMATION

### Supplementary notes

Detailed descriptions for cases I2-I7 and additional methods, results and figures.

### Supplementary File 1

Excel file containing the worksheets “summary”, “gene”, “domains”, “CNVs_cases”, “CNV_genes”, “variants_cases”, “all_missense”, “all_AAdel”, “variants_annotated”, “variants_reviewed” and “publications_reviewed” used for Fig. 2. The “summary” worksheet contains detailed descriptions of all worksheets and the respective data columns.

### Supplementary File 2

Excel file containing the worksheets “summary”, “CP02-Panel_Genes”, “CP02-Panel_Design”, “HNF1B_primers”, “PKD1_LR-PCR” and “PKD1_Seq” describing the targeted panel design and primers used for additional sequencing. The “summary” worksheet contains a detailed description of all other worksheets and the respective data columns.

