## Supplementary Notes for "Prenatal diagnosis of *HNF1B*-associated renal cysts: Need to differentiate intragenic variants from 17q12 microdeletion syndrome?"

### SUPPLEMENTARY CASE REPORTS

#### Detailed clinical description of individuals with 17q12 microdeletion syndrome

##### Individual 2 (I2)

Individual I2 is a male individual, born at 41 weeks of gestation by Caesarean section. Pregnancy history involved gestosis. Birth parameters were in normal ranges. After birth, a ventricle septum defect was diagnosed, not requiring surgical repair. He presented at the age of 8 years and 8 months with developmental delay. He could walk independently at the age of 13 months and spoke first words at age 12 months. For several years he received physiotherapy, occupational and speech therapy. Although he was initially enrolled in a regular school, he changed to a school for children with special needs due to learning difficulties. He also displayed oppositional behaviour. At the time of last assessment, height was 130.3 cm (25. P) and weight 29.2 kg (50-75. P). Head circumference was 56.5 cm (97. P). Minor dysmorphic features included a long face, a high nasal root, a wide nasal bridge, prominent front teeth, a deep philtrum, high palate, auricular tags, short fingers, broad thumbs and big toes. Cranial magnetic resonance imaging (cMRI) did not show any abnormalities. Up to the point of the last clinical assessment kidney evaluation was not performed. Information about liver, pancreas and urogenital anomalies as well as diabetes, abnormal levels of liver enzymes, hypomagnesemia and hyperuricemia was not available. Ophthalmologic anomalies were not reported. Individual I2 was the second child of non-consanguineous healthy parents.

##### Individual 3 (I3)

The mother of individual I3 reported premature contractions during pregnancy. The boy was born at 41 weeks of gestation by Caesarean section with normal birth parameters. Motor and speech milestones were at first age-appropriate. However, hyperactivity and restlessness were observed by the parents already from 12 months on. He was also very focused on a well-regulated daily routine, control and order and exhibited abnormal social behaviour. At the age of 12 he attended a special school and was referred for genetic testing due to learning difficulties and behavioural abnormalities. The physical examination showed a weight of 31.4 kg (10. P), a height of 141 cm (10. P) and a head circumference of 52 cm (3-10. P). He exhibited hair swirls on the forehead, a high and narrow palate, relatively large incisors, mild retrognathia, a single transverse palmar crease on the right hand, short fingers and broad thumbs. Renal examination was not performed at that time. On reassessment at 15 years and 8 months of age, I3 was diagnosed with Asperger syndrome. Testing revealed mild ID (IQ of 63). cMRI was not performed and eye abnormalities were not registered.

Sonographic examination of the kidneys at that time showed bilateral renal cysts, whereas liver and pancreas were not examined. Serum creatinin was 0.87 mg/dl, urine total protein 247mg/l, albumin/creatinine ratio 179 mg/g, but hepatic transaminases and magnesium levels were normal. Plasma uric acid concentration was not measured. No sign of diabetes was detected. The parents of individual I3 were healthy and non-consanguineous. They experienced a miscarriage at 7 weeks of gestation. His older sister exhibited hemiplegia due to a brain haemorrhage, and a maternal half-brother was diagnosed with ADHS.

**Individual 4 (I4)**

Individual I4 was a 6 year old female, born at 38 weeks of gestation. During pregnancy, gestosis was diagnosed in the mother. Birth parameters were normal. She was referred to our genetic clinics because of motor developmental delay and short stature. She was able to walk independently after the age of 18 months; physiotherapy and occupational therapy were required. Because of learning and concentration difficulties, she enrolled in a special school. I4 had increased levels of liver enzymes (GOT: 76.3 U/l, GPT: 59.6 U/l and GGT: 27.6 U/l), a sonographic examination of the liver though was not performed. Pancreas und urogenital tract examination was also not available. Her height was measured at 107.9 cm (< 3. P), she showed though good response to growth hormone treatment as indicated by the increase of the predicted final adult height. Weight was 16.7 kg (3. P) and head circumference 53 cm (75. P). Subtle dysmorphic features included medial flared eyebrows, long eyelashes, broad and flat nose root, thin upper lip and hypertrichosis. At age of 12 years I4 was reported to additionally present with autistic-like behaviour. Her IQ was low-normal (IQ of 80) and she was attending a regular school. cMRI revealed enlarged brain ventricles. Furthermore, I4 was diagnosed with Brown´s syndrome (rare form of strabism). Renal abnormalities or diabetes have not been detected to date. Individual I4 was not examined regarding hypomagnesemia and hyperuricemia. The parents of individual I4 were healthy and non-consanguineous. A cousin of the mother exhibited a global developmental delay.

**Individual 5 (I5)**

Female individual I5 was hospitalized due to polydipsia and fatigue at 18 years of age. She had been diagnosed with diabetes mellitus at the age of 14. After further investigation multiple bilateral renal cysts were detected, which 4 years later resulted in stage II of chronic kidney disease. Sonographic evaluation of liver and pancreas and examination of urogenital tract were not performed. Plasma uric acid levels were increased, liver transaminases were normal and plasma magnesium concentration was not measured. Apart from a mild strabism she did not show any other ocular anomalies. At the time of the evaluation weight was 48.7 kg (25. P) and height 165.5 cm (50-70. P). She had no psychomotor impairment and attended a regular school. Individual I5 was the second child of reported non-consanguineous parents. Her father as well as the brother and mother of her father were likewise diagnosed with renal cysts. Nevertheless, none of them was a carrier of the 17q12 microdeletion identified in I5. Therefore, the presence of an additional polycystic kidney disease in this family was discussed, but not followed up further.

**Individual 6 (I6)**

Individual I6 presented at age 28 years. Due to motor developmental delay during early childhood supporting measures had taken. Speech and cognitive development were normal, and learning difficulties or behavioral abnormalities were never noted. She reported recurrent urinary tract infections in childhood and puberty. At age 24 years she experienced a single seizure. In the same year primary biliary cirrhosis due to liver cysts and elevated levels of hepatic enzymes (GOT: 69 U/l, GPT: 121 U/l and GGT: 109 U/l) were diagnosed. Information about plasma magnesium and uric acid levels was not available. Blood sugar levels were repeatedly increased. After the age of 26 and, while she was pregnant, bilateral renal cysts and end-stage renal disease of the right kidney, were identified. Her weight was 48.4 kg (3-10. P) and her height 156.5 cm (3-10. P). Head circumference was 57.5 cm (90-97. P). Individual 6 did not exhibit any facial or physical dysmorphic features. Ophthalmic examination and cMRI were not performed. Her parents were non-consanguineous. Her father was diagnosed with epilepsy and diabetes type 1, whereas her paternal grandfather had diabetes type 2.

**Individual 7 (I7)**

Individual I7 was referred to genetic testing at age 45 years due to an end-stage renal disease requiring dialysis. At the age of 4 months she was diagnosed with right kidney aplasia and left kidney hypoplasia. She also experienced recurrent interstitial nephritis of the left kidney. Two kidney transplantations, the first at age 19 years and the second at age 36 years were unsuccessful. Data regarding sonographic examination of the liver or pancreas and determination of plasma magnesium and uric acid concentration was not available. Diabetes mellitus occurred after the second transplantation and was attributed to immunosuppressive therapy. Furthermore, gynecological examination revealed uterus aplasia, but normal ovaries. At the age of 36 years, cysts were removed from the left ovary. Ureters were structurally normal. Strabism convergens, aortic insufficiency and joint hyperextension were also reported. Individual 7 showed no neurological deficits or psychomotor delay. She was the third child of non-consanguineous parents. Both parents and siblings had a normal renal function.

**Tab. S1**: Summary of 12 clinical features in herein described individuals with *HNF1B* aberrations using Human Phenotype Ontology (HPO) terms. The following abbreviations and symbols are used: HP:, Human Phenotype; +, present; -, absent; NA, not analyzed

| | **Individual 1** | **Individual 2** | **Individual 3** | **Individual 4** | **Individual 5** | **Individual 6** | **Individual 7** |
|---|---|---|---|---|---|---|---|
| HP:0000077 | + | - | + | - | + | + | + |
| HP:0012758 | - | + | + | + | - | + | - |
| HP:0000819 | +? | NA | - | - | + | + | - |
| HP:0000078 | - | NA | - | NA | - | - | + |
| HP:0002910 | - | NA | NA | + | NA | + | NA |
| HP:0410042 | - | NA | NA | NA | - | + | - |
| HP:0001732 | - | NA | - | NA | - | - | - |
| HP:0012443 | NA | - | NA | + | NA | NA | NA |
| HP:0004322 | - | - | - | + | - | - | - |
| HP:0000478 | - | - | - | + | + | NA | + |
| HP:0004921 | - | NA | - | NA | NA | NA | NA |
| HP:0002149 | - | NA | NA | NA | + | NA | NA |

**Description of additional CNVs identified in affected individuals**

Microarray analysis revealed in individual I6 an additional 319 kb microduplication on X chromosome (arr[GRCh37] Xq27.1(139902009_140220878)x3) encompassing four genes; *MIR320D2, SPANXB2, SPANXF1* without OMIM phenotype, and *SPANXB1*. The latter belongs to *SPANX* gene cluster on chromosome X, yet its function remains to be determined. No association with either developmental delay or renal disease has been reported, to date. Furthermore, the same duplication was previously identified in an individual of our control cohort, and several smaller but overlapping aberrations have been reported in control population. Screening of the parents for this aberration was not performed. We classified it as a CNV of unknown significance (VUS), but given that it is a gain, we consider a clinical relevance unlikely.

### SUPPLEMENTARY METHODS

#### Panel sequencing and variant confirmation

For I1 sequencing was performed on an Illumina MiSeq system using a customized TruSight Sequencing Panel (Illumina Inc., San Diego, USA). Sequencing reads were aligned and processed as previously described.[1] Four genes, *PKD1* (NM_001009944.2), *PKD2* (NM_000297.3), *PKHD1* (NM_138694.5) and *HNF1B* (NM_000458.3) were analyzed using the SeqNext module of the SeqPilot software (JSI medical systems, Ettenheim, Germany) (also see Supplementary File 2, sheets "CP02-Panel_Genes" and "CP02-Panel_Design").

#### Sanger sequencing of *HNF1B*

The *HNF1B* variant identified in I1 and segregation of this variant in the family was confirmed by Sanger sequencing using standard procedures and the following primers: "HNF1B_Exon3": 5′-TTGCAAAGCTTAGTTAGACGAGG-3′ and 5′-AACTAGTGTCTCAATATCCCAGGAC-3′ (also see Supplementary File 2, sheet "HNF1B_primers").

#### Long Range PCR and Sanger sequencing of *PKD1*

Due to the complex nature of the large *PKD1* gene (46 exons) locus, with high GC content and high homology of the first 33 exons with six pseudogenes which arised through duplication of chromosome 16, we additionally screen *PKD1* using Long-Range PCR (LR-PCR), if no variant through high-throughput short-read sequencing is identified. The protocol used is based on the method described by Tan et al. in which 9 amplicons are amplified by LR-PCR and then directly Sanger sequenced using internal primers.[2] Primer sequences for LR-PCR and Sanger sequencing can be found in Supplementary File 2, sheets "PKD1_LR-PCR" and "PKD1_Seq", respectively. We re-designed 11/44 internal sequencing pairs. Our protocol uses the Expand Long Range dNTPack (Sigma-Aldrich, St. Louis, Missouri, USA) with 40 ng of genomic DNA in a volume of 25 µl for LR-PCR, Ampure beads (Beckman Coulter, Brea, Orange County, USA) for PCR purification and standard procedures for Sanger-sequencing.

#### Chromosomal microarray analysis

Chromosomal microarray analysis (CMA) for individuals I2, I3 and I6 was performed with a Genechip 6.0 Mapping SNP-Array, for I4 and I5 with a CytoScan HD-Array (both: Affymetrix, Santa Clara, USA) and for I7 with an Oligo-Microarray Human Genome 244A (Agilent Technologies, Santa Clara, USA) according to the manufacturer's instructions. Copy number variants (CNV) were analyzed using the Affymetrix Chromosome Analysis Suite 3.1.0.15

(ChAS; Affymetrix, Santa Clara, USA) software against an database of 3,500 in-house samples and the Database of Genomic Variants (DGV) for aberrations sizing ≥ 100 kb. FISH with a locus specific DNA probe RP11-27K22 (in-house) or MLPA analysis with the Kit P297 (MRC Holland, Amsterdam, The Netherlands) were used for confirmation of the 17q12 microdeletions in affected individuals and for segregation testing in the parents.

**Collection and standardization of *HNF1B* variants reported as pathogenic**

To analyze potential clustering and important domains we intended to generate an up-to-date list of intragenic *HNF1B* variants. By searching PubMed using the terms "HNF1B" or alternatively "TCF2" together with the terms "mutation" or "variant" and additionally searching through the identified articles citations, we identified 88 publications describing intragenic variants in *HNF1B* in individuals presenting with *HNF1B*-associated phenotypes. We did not include the whole gene deletions additionally described in some of these publications, as the breakpoints were often not described, and our intention was to analyze intragenic variants. We also included (likely) pathogenic variants from the ClinVar[3] and LOVD[4] databases. Variants were harmonized to the NM_000458.3 transcript and the hg19 reference genome based on Human Genome Variation Society (HGVS) recommendations using VariantValidator.[5] As most reported variants had not been scored using a 5-tier classification system, we applied the American College of Medical Genetics and Genomics (ACMG) criteria[6] using InterVar[7] and additionally manual curation. Such identified variants were annotated with computational scores and databases from dbNSFP[8] version 2.9.3 and variant frequencies from the gnomAD database using SnpEff/SnpSift.[9, 10] Additionally, we annotated CADD v1.4 PHRED scores using the online annotation service.[11] To visualize the variability of 17q12 microdeletions we downloaded the genomic coordinates of 33 individuals with CNVs in this region described as (likely) pathogenic in the DECIPHER[12] database. All variant information is provided in Supplementary File 1 and the standardized intragenic variants have been submitted to both the ClinVar and LOVD databases.

**Review of reported clinical features in literature**

To analyze the variable reporting of clinical symptoms in the literature we seached all 88 above identified publications for 12 features previously associated with *HNF1B*-disorders. If the presence or absence of the respective feature was mentioned we scored the publication as "1", if not as "0". Additionally, we categorized the publication type (review, case reports (<3 cases), case series (≥3), screening of *HNF1B* and/or other genes in larger cohorts), the medical specialty (according to the journal topic or first/last authors affiliations), the *HNF1B* variant types analyzed (17q12del and/or SNV/indel) and whether the publication described born individuals or fetal cases. The detailed results of this review are represented in (Supplementary File 1, sheet “publications_reviewed”) and summarized in Tab. 2.

**Spatial clustering analysis of pathogenic variants**

To analyze a potential clustering of variants in the domains of the linear protein representation, we generated density plots of (likely) pathogenic missense variants and likely truncating variants with the “geom_density” function (“adjust” parameter set to 1/2) in ggplot2 and calculated absolute and local maxima (Fig. 2B). Further we calculated empiric distributions for drawing 75 missense variants from all possible 3,687 missense substitutions and estimated p-values for the observed distribution of the (likely) pathogenic missense variants described in literature and databases (Fig. S2).

**Computational analysis of single amino-acid deletions and missense variants**

We generated all possible missense variants and 3 base pair deletions in the *HNF1B* gene region of the hg19 reference (chr17[hg19]:36044434-36107096) as variant call format (VCF) files which were annotated as described above. Missense variants and single amino acid (AA) deletions affecting the NM_000458.3 transcript of *HNF1B*, were filtered for further analyses. Variants, which were additionally annotated as potentially affecting splicing or deleting AAs while generating a new codon (“delins”), were excluded to generate the possible 3 bp-deletion variants leading to the deletion of a single AA (“AAdel”) (Supplementary File 1, sheets “all_missense” and “all_AAdel”). We used this data to analyze protein regions of higher conservation by plotting all missense variants sorted by AA position with their respective CADD score and fitted a generalized additive model (Fig. 2B, lower panel). To analyze the conservation of HNF1B protein regions we compared the scores of all possible missense variants in the respective domains (Fig. S1).

**Protein structure analysis of the Gly239del *HNF1B* variant**

We analyzed the spatial proximity of the Gly239 AA position to the DNA double-helix and AA positions important for DNA binding using Pymol (Version 1.8.6.0; Schrödinger, LLC) with the tertiary protein structure data of HNF1B bound to DNA (PDB-ID: 2H8R[13]) (Fig. 3).

**Data analysis and plotting**

The variant data provided in Supplementary File 1 (Excel; Microsoft Corporation, Redmond, USA) was analyzed and plotted using R language version 3.5.1 with RStudio IDE version 1.1.463 (RStudio, Inc.) with packages from the tidyverse collection. Libraries "broom", "cowplot", "fuzzyjoin", "ggrepel", "ggsignif", "plyr", "readxl", "Rmisc", "tidyverse", “GenomicRanges”, “Gviz”, “trackViewer” and “ggplotify”. Illustrator / Photoshop CC 2018 (both: Adobe Systems, San José, USA) or Inkscape 0.92.3 (https://inkscape.org/) were used to adjust main Fig. 2 for parts which could not be directly composed in R and to compose main Fig. 1 and Fig. 3 from primary image data.

**SUPPLEMENTARY FIGURES**

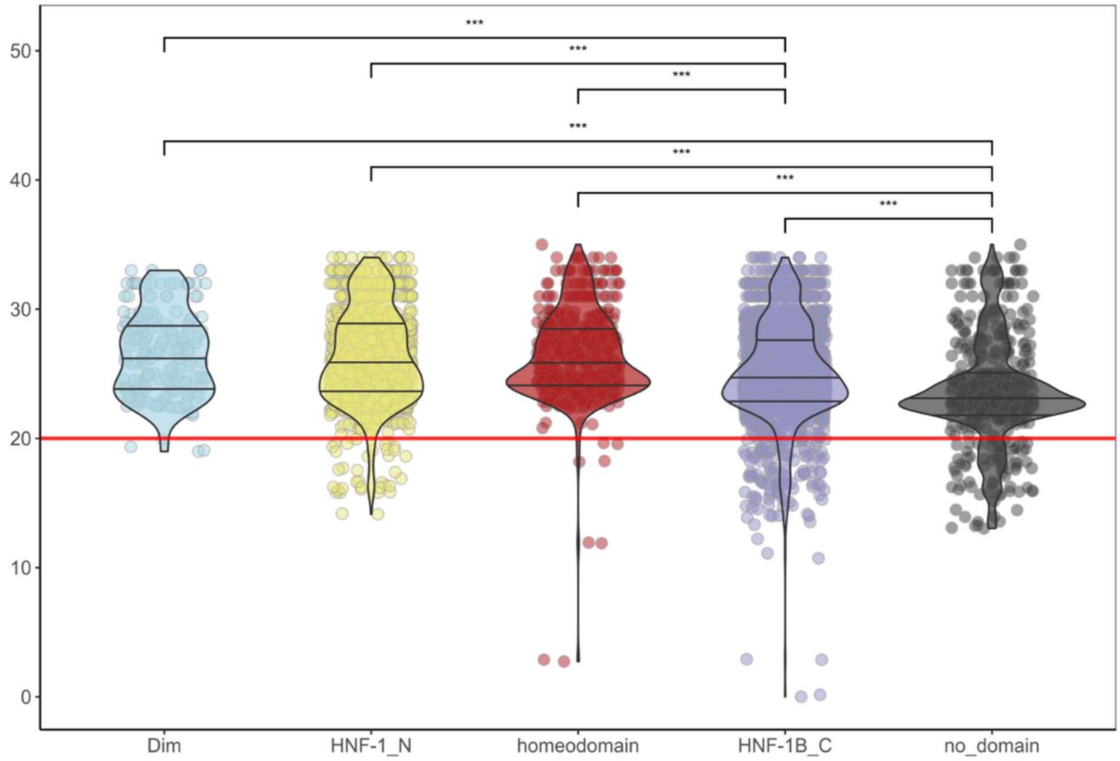


**Fig. S1 | comparison of CADD scores for missense variants in HNF1B domains**

Violin- and scatter-plot comparing the CADD scores[11, 14] for all possible missense variants (Supplementary File 1, sheet "all_missense") in known domains of the HNF1B protein (Dim, N-terminal dimerization domain: light blue; HNF-1_N, N-terminal domain: yellow; homeodomain, DNA binding domain: red; HNF-1B_C, C-terminal transactivation domain: blue) and protein regions without domain assignment ("no_domain": grey). The CADD scores in all 4 domains are significantly higher than in the non-domain regions. Additionally, the CADD scores are higher in the two N-terminal domains and the homeodomain, required for dimerization with HNF1A and DNA binding, than in the HNF-1B_C domain. This observation agrees with the reported clustering of identified missense variants in the homedomain and the second half of the HNF-1_N domain (Fig. 1, Fig. S2). Two-sided Wilcoxon signed-rank used for significant testing. ***: P<0.001.

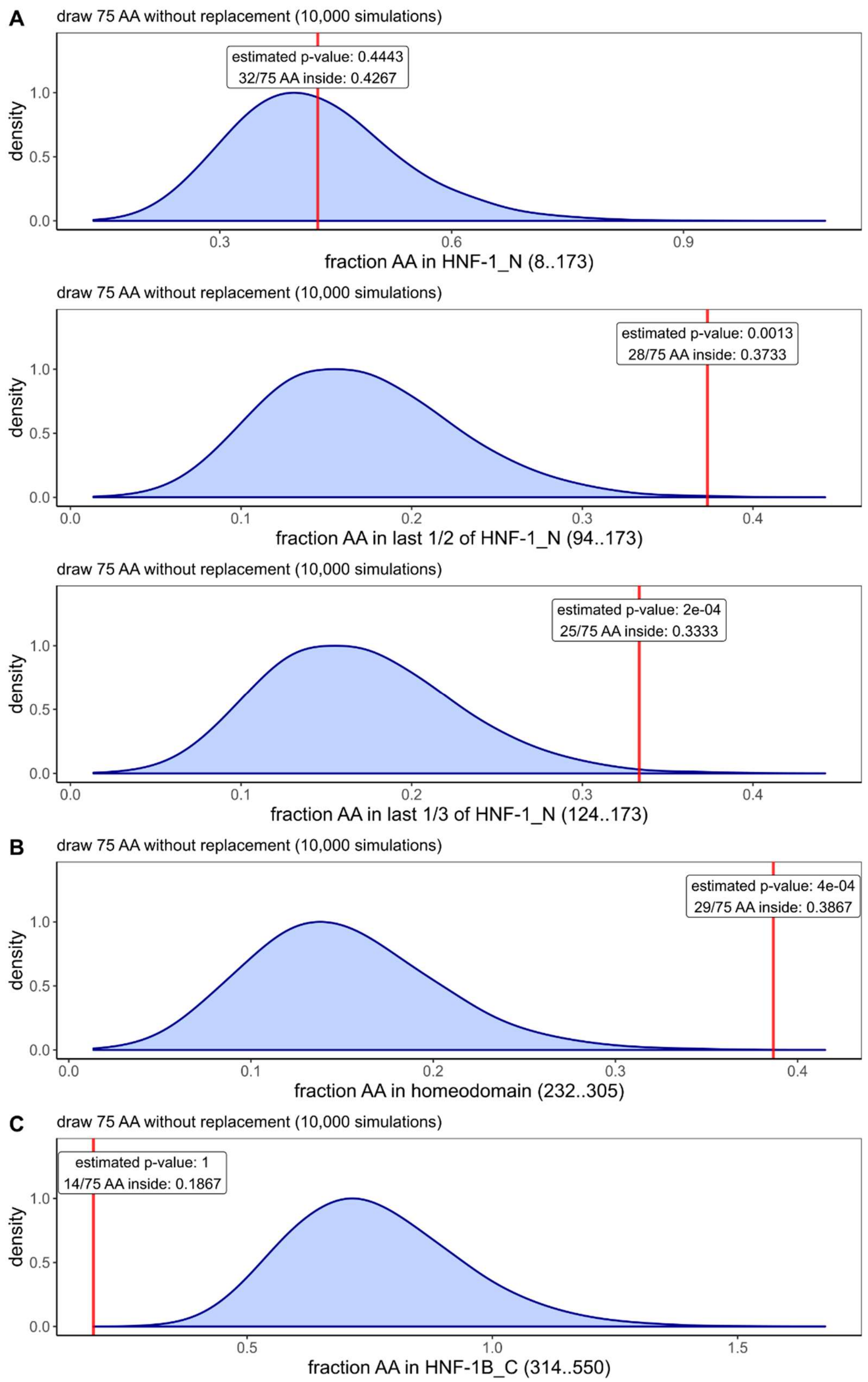


**Fig. S2 | Analysis of the variant clusters in the HNF-1_N and homedomain domain**

Empiric distribution for drawing 75 unique (e.g. number of (likely) pathogenic from literature in Fig. 2B) missense variants from all possible 3,687 missense substitutions (Supplementary

File 1 sheet “all_missense”) in the HNF-1_N (A), homedomain (B) and HNF-1B_C (C) domain of HNF1B without replacement. While the 32 missense variants in the complete HNF-1_N domain (AAs 8-173) indicate no enrichment (A: upper panel), the enrichment in the second half (A: middle panel; AAs 94-173; containing 28 described missense variants; estimated p-value = (draws ≥ 28/75 in) + 1) / (all draws + 1) ~ 0.0013) and the last third (A: lower panel; AAs 124-173; containing 25 described missense variants; estimated p-value = (draws ≥ 28/75 in) + 1) / (all draws + 1) ~ 2e-4) of this domain is highly unlikely by chance assuming uniform mutation distribution. Also, the enrichment in the homedomain (B; AAs 232-305; containing 29 described missense variants; estimated p-value = (draws ≥ 29/75 in) + 1) / (all draws + 1) ~ 4e-4) is highly unlikely. The HNF-1B_C domain (C; AAs 314-550; containing 14 described missense variants) in contrast shows no enrichment of described missense variants (estimated p-value = (draws ≥ 14/75 in) + 1) / (all draws + 1) ~ 1). In summary, this analysis confirms the clustering of pathogenic missense variants in the homedomain and the second half of the HNF-1B_C which are both important for DNA binding.
